# Residual Sanitization of Three Human Respiratory Viruses on a Hard, Non-Porous Surface

**DOI:** 10.1101/2023.03.02.530883

**Authors:** Luisa A. Ikner, Andrew B. Rabe, Charles P. Gerba

**Author notes:** Contact information for the corresponding author: Luisa A. Ikner, PhD, Assistant Professor, Department of Environmental Science, Water & Energy Sustainable Technology (WEST) Center, Tucson, AZ 85745.

## Abstract

Human pathogenic viruses that are present in bioaerosols released by coughing, sneezing, or breathing can contaminate fomites and other inanimate environmental surfaces. Most are enveloped respiratory viruses that are vulnerable to inactivation by a broad spectrum of antimicrobial actives. Quaternary ammonium compounds are highly diverse in structure and are among the most widely utilized antimicrobial agents. The objective of this study was to evaluate two commercially available, ready-to-use quaternary ammonium compound-based disinfectants (one of which also contains a surface binding agent) for antiviral activity against Influenza A (H1N1), human coronavirus 229E, and SARS-CoV-2 (Washington) following a rigorous procedure of wear and abrasions with regular re-inoculations of virus in the presence of a 6% organic soil load. Formulation TF-A demonstrated variable residual efficacy against the three viruses, achieving log_10_reductions of 1.62, 3.33, and 0.92, respectively. Formulation TF-B lowered each test virus by greater than 3-log_10_to non-detectable levels on all carriers in demonstration of residual antiviral activity.

## Introduction

Communicable human respiratory viruses including influenza, coronavirus, rhinovirus, and respiratory syncytial virus (RSV) co-circulate seasonally. Illness from such viral pathogens result in increased rates of morbidity and school absenteeism, and contribute to losses in productivity (Papadopoulos, 2017; Donaldson et al., 2021; Macias et al., 2021) The advent of COVID-19 introduced SARS-CoV-2 into the pool of potential respiratory pathogens of risk to humans and prompted wide-scale mitigation efforts including masking, social distancing, enhanced hand hygiene, and intensive disinfection protocols. Inhalation of infectious airborne aerosols has been proposed as the primary mode of transmission for enveloped respiratory pathogens such as influenza (Cowling et al., 2013) and SARS-CoV-2 (Tang et al., 2020). Virus survival within bioaerosols is impacted by relative humidity, absolute humidity, temperature (Prussin et al., 2018), and the cumulative exposure of solutes that increase in concentration as aerosols evaporate (Lin & Marr, 2020). Following deposition onto hard, non-porous surfaces, infectious SARS-CoV-2 can remain viable for several days under low temperature and low humidity regimes (Aboubakr et al. 2020), while Influenza A (H1N1) can remain infectious on stainless steel for up to 24 hours (Oxford et al., 2014).

Disinfection is an effective strategy to alleviate levels of pathogenic viruses deposited onto hard, nonporous surfaces, thereby lowering the risk of transmission via fomites. The susceptibility of viruses on surfaces to inactivation by ready-to-use liquid disinfectants (delivered by spray device or other means) is largely related to structure, with enveloped viruses such as influenza A, human coronavirus 229E (HCoV-229E), and SARS-CoV-2 being more susceptible to a broader variety of chemistries including surfactants, quaternary ammonium compounds, and alcohols than non-enveloped viruses (Wolff et al., 2005). Standard ready-to-use disinfectants are formulated to inactivate microorganisms within the contact times specified on the product label and generally do not provide for residual antimicrobial activity, especially when wiped dry. Surface-anchored quaternary silanes have offered an alternative to impart residual self-sanitizing properties to treated surfaces. A C18 quaternary silane was recently evaluated against SARS-CoV-2 and remained effective for 47 days after the product was applied (Caschera et al., 2022). However, the quaternary silane-treated carriers remained undisturbed during the 47-day period, whereas synthetic environmental surfaces within built environments are likely to be subjected to periodic cleaning and/or disinfection practices as well as recontamination over time due to touching or other types of deposition events. Disinfectants that are comprised in part of surface anchoring or binding agents and claim residual antimicrobial effectiveness should be subjected to more rigorous evaluations that incorporate elements of friction and recontamination to ensure that such claims are valid.

The objective of this study was to compare the residual self-sanitizing properties of two commercially available quaternary ammonium compound-based spray disinfectants - one bearing a United States Environmental Protection Agency (US EPA) residual kill label claim of 99.9% against bacteria and containing a surface binding chemistry (polyethyloxazoline) - using a wear and abrasion procedure with repeated virus inoculations in the presence of 6% organic soil against three prominent respiratory pathogens of concern: Influenza A (H1N1); human coronavirus 229E, and SARS-CoV-2 (Washington isolate).

## Materials and Methods

### Viruses and Cell Lines

Human coronavirus 229E (HCoV-229E; VR-740) was procured from the American Type Culture Collection (ATCC; Manassas, VA, US) and propagated using the MRC-5 cell line (ATCC CCL-171). SARS-CoV-2 (Washington strain) and an isolate of Influenza A catalogued during the 2009 H1N1 Pandemic were obtained through BEI Resources, NIAID, and NIH, with designations of SARS-Related Coronavirus 2, Isolate USA-WA1/2020 (NR-52281), and Influenza A Virus, A/New York/18/2009 (H1N1)pdm09 (Tissue Culture Adapted; NR-15268), respectively. Stocks of SARS-CoV-2 were prepared within a Biosafety Level 3 laboratory suite using the Vero E6 (ATCC CRL-1586) cell line. Madin-Darby Canine Kidney (MDCK) cells were used to propagate stocks of Influenza A (H1N1)pdm09 for the study. HCoV-229E and SARS-CoV-2 propagations were performed when the MRC-5 and Vero E6 host cell monolayers reached 80% to 90% confluency, and Influenza A (H1N1) infections were performed when MDCK monolayers were 90% to 95% confluent. Flasks were infected at a multiplicity of infection (moi) range of 0.1 to 0.01 and incubated for 30 minutes within a 5% CO_2_atmosphere at the appropriate temperature [37°C: SARS-CoV-2; 35°C: HCoV-229E, and 33°C: Influenza A (H1N1)pdm09]. Maintenance medium was then added to the infected flasks [2% FBS Eagle’s Minimal Essential Medium (MEM) for SARS-CoV-2 and HCoV-229E, and MEM supplemented with 10 μg/mL TPCK-treated trypsin, 10mM of HEPES buffer, and 0.125% (w/v) bovine serum albumin, termed as Influenza Infection Medium, for Influenza A (H1N1)pdm09]. After 48 to 72 hours, infected flasks were subjected to two freeze-thaw cycles and the lysates were centrifuged for 10 minutes (1,000 x g) to pellet the cell debris. Viruses in the supernatant were further concentrated overnight at 2-5°C with stirring using polyethylene glycol [8000 g/mol; 13% (w/v)] and 0.5M NaCl. Virus flocs were then pelleted by centrifugation (10,000 x g, 30 minutes) and resuspended using Dulbecco’s Phosphate-Buffered Saline (DPBS) in 3% to 10% of the original lysate volume. Virus stocks were aliquoted and stored at -80°C. Prior to use in the study, stocks were titered by dilution (1:10) and plating onto their respective host cell monolayers plated in multi-well trays using the tissue culture infectivity dose at the 50% endpoint (TCID_50_) technique. Viral stock titers were calculated using the Spearman-Karber algorithm as described by Hierholzer and Killington (1996).

### Preparation of Carriers

The steps and time-range requirements of the wear and abrasion test procedure are shown in Table 1. Glass carriers (1-in^2^) were pre-sanitized using 95% ethanol and wiped dry within a biosafety cabinet. On the day prior to initiation of the wear-testing, two commercially available disinfectant formulations and the control substance [sterile deionized water (DI-H_2_O)] were applied to the carriers via spray device within an environmental chamber maintained at 20°C to 23°C (45% to 48% relative humidity. The commercial test formulations evaluated in the current study were designated as Test Formulation A (TF-A) and Test Formulation B (TF-B), and their respective active ingredients are listed in Table 2. Each test formulation was applied using the spray device and nozzle supplied by the manufacturer. Carriers were sprayed from 6 to 8 inches in distance at an angle of 45°. Additional carriers were treated using sterile DI-H_2_O delivered via Preval sprayer at the same distance and spray angle specified for the test formulations to assess virus inactivation in the presence of an inert chemistry. Five spray coatings of the test and control substances were applied per carrier. One set of DI-H_2_O control carriers (n=4) underwent wear and abrasion testing; the other DI-H_2_O control set (n=4) was not subjected to wear-testing. The treated test and control carriers were cured overnight within the temperature range of 20-23°C (45-48% relative humidity).

**Table 1.**
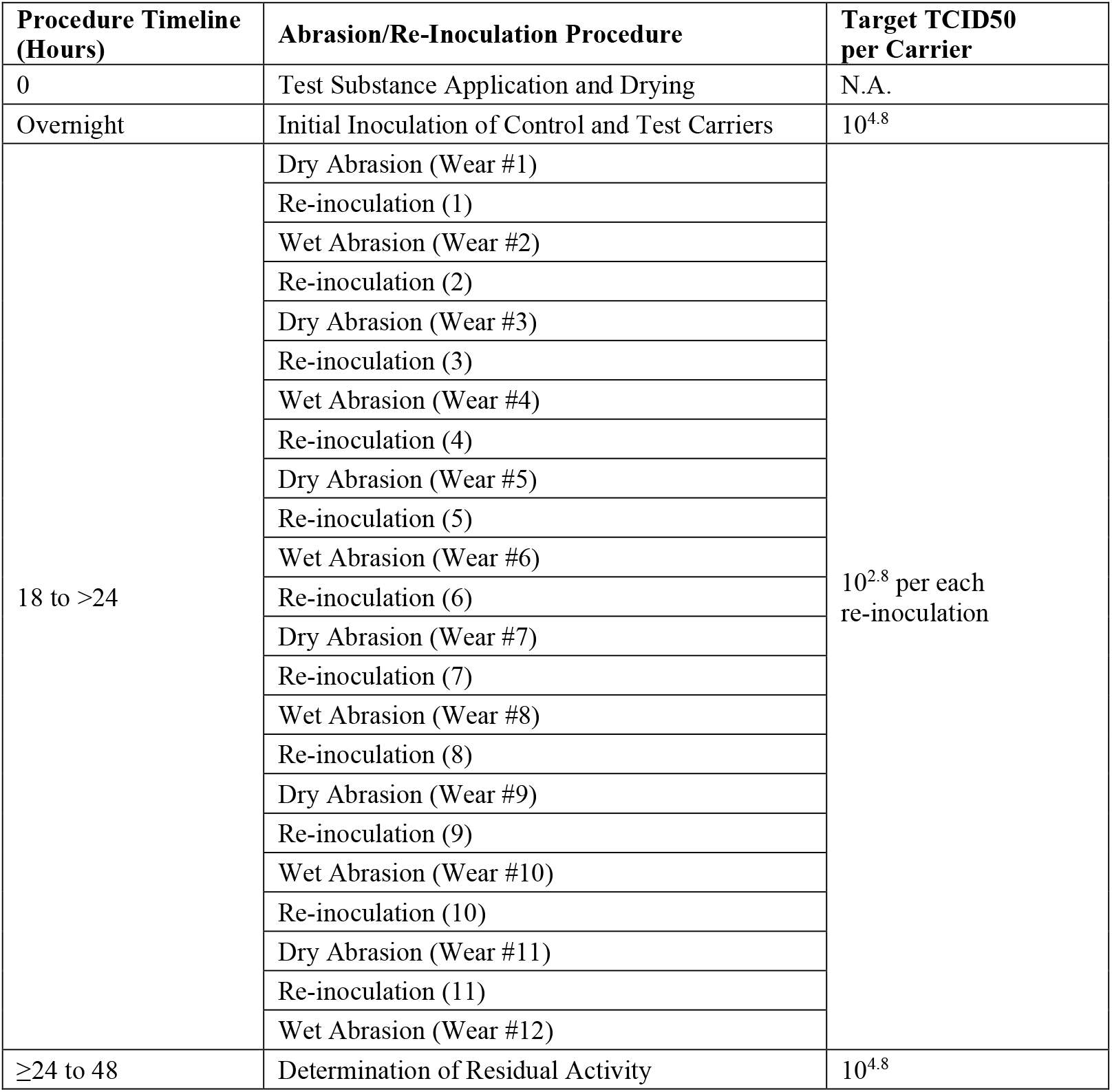
Wear and Abrasion Test Procedure with Reinoculation Regimen

| Procedure Timeline (Hours) | Abrasion/Re-Inoculation Procedure | Target TCID<sub>50</sub> per Carrier |
| --- | --- | --- |
| 0 | Test Substance Application and Drying | N.A. |
| Overnight | Initial Inoculation of Control and Test Carriers | $10^{4.8}$ |
| 18 to >24 | Dry Abrasion (Wear #1) | $10^{2.8}$ per each re-inoculation |
|  | Re-inoculation (1) |  |
|  | Wet Abrasion (Wear #2) |  |
|  | Re-inoculation (2) |  |
|  | Dry Abrasion (Wear #3) |  |
|  | Re-inoculation (3) |  |
|  | Wet Abrasion (Wear #4) |  |
|  | Re-inoculation (4) |  |
|  | Dry Abrasion (Wear #5) |  |
|  | Re-inoculation (5) |  |
|  | Wet Abrasion (Wear #6) |  |
|  | Re-inoculation (6) |  |
|  | Dry Abrasion (Wear #7) |  |
|  | Re-inoculation (7) |  |
|  | Wet Abrasion (Wear #8) |  |
|  | Re-inoculation (8) |  |
|  | Dry Abrasion (Wear #9) |  |
|  | Re-inoculation (9) |  |
|  | Wet Abrasion (Wear #10) |  |
|  | Re-inoculation (10) |  |
|  | Dry Abrasion (Wear #11) |  |
|  | Re-inoculation (11) |  |
|  | Wet Abrasion (Wear #12) |  |
| ≥24 to 48 | Determination of Residual Activity | $10^{4.8}$ |

**Table 2.**
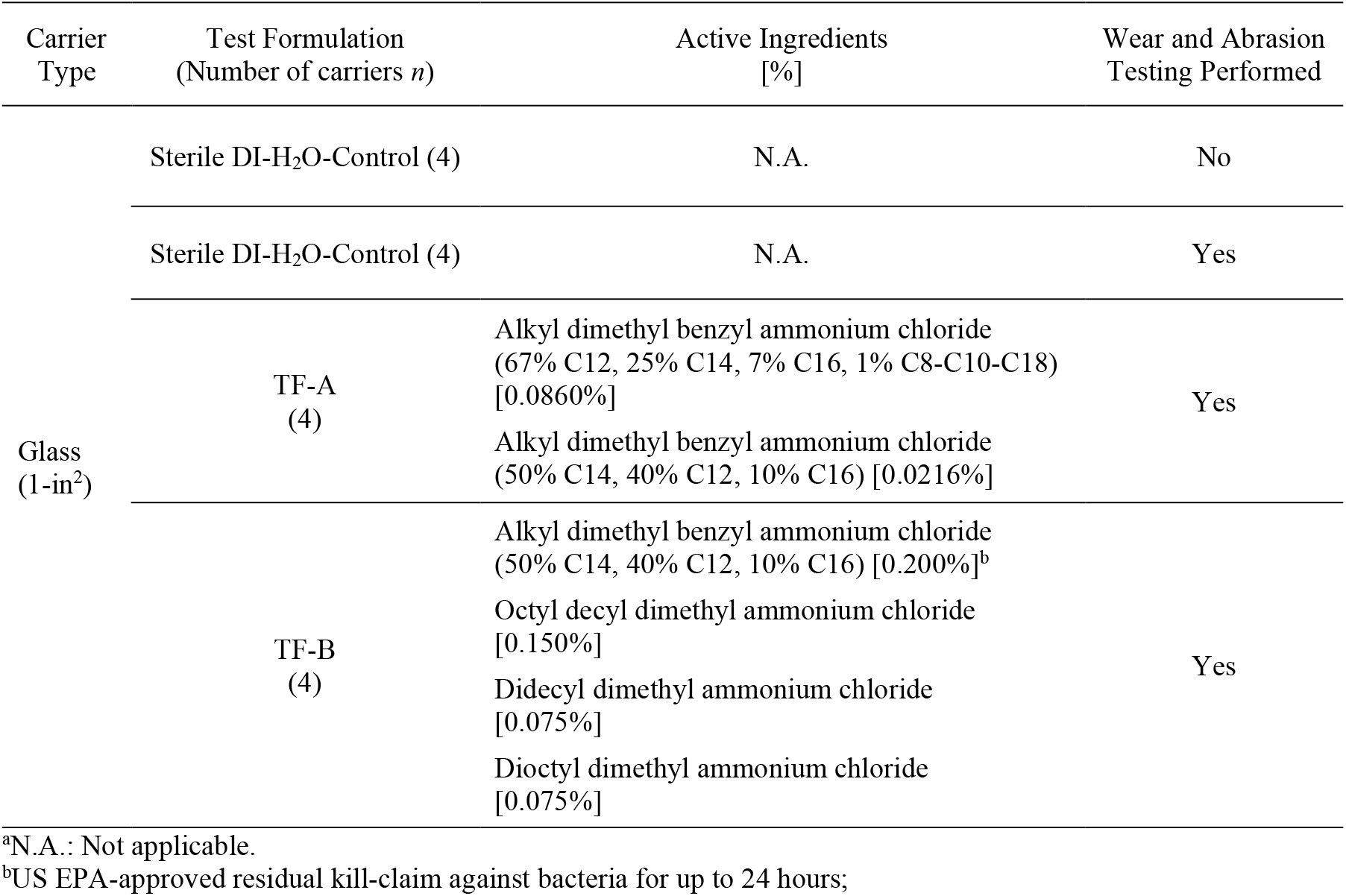
Description of Test Formulations and Prepared Control and Test Carrier Designations

On the next day, four control and five four carriers per formulation were placed into sterile petri dishes containing one layer of Whatman #42 filter paper. Only carriers demonstrating complete (100%) coverage of the applied formulations were used for the wear and abrasion testing. One additional carrier per control and test formulation was prepared to assess cytotoxicity on host cells and to validate test substance neutralization.

### Initial Virus Inoculation followed by the Wear and Abrasion Test Procedure

Virus stocks were thawed just prior to testing and amended with fetal bovine serum to obtain a 6% (v/v) organic soil load. Each control and test carrier deemed suitable for experimentation was inoculated with 10 μL of the test virus at a minimum recoverable target density of 4.8-log_10_TCID_50_per carrier (Table 1). The inoculum was spread to within 1/8” of the carrier edge using a bent pipette tip. The carriers were then incubated at 37°C for 30 to 35 minutes with the petri dish lids slightly ajar until complete drying was observed.

Cotton wipers (TexWipe #TX309, MTE Solutions, Kernersville, NC, USA) cut into dimension of 9” x 4” along the thread pattern were pre-sterilized by autoclaving for 15 minutes. Foam liners (FoamWipe #TX704, MTE Solutions) were also prepared prior to wear testing by cutting the material into 4” x 2” pieces. The Gardco® Washability & Wear Tester (Model #D10V, Paul M. Gardco Co. Inc., Pompano Beach, FL, USA) was placed into a Class II A2 biological safety cabinet (Class II B2 for SARS-CoV-2) and decontaminated using 95% ethanol. Two Gardco® abrasion boat weights (Model #13931) were fitted with one foam liner and one cotton wiper each and verified to be 1,084 ± 1.0g prior to use. The carriers were placed in singlet onto the abrasion test surface and then subjected to a complete wear and abrasion cycle consisting of four total passes (one pass to the left, one pass to the right, followed by one pass again to the left and then one to the right) at a speed of 2.25-2.5 for a total surface contact time of 8 to 10 seconds. This first wear exposure series was considered a “dry” abrasion cycle. Upon completion of the four passes, the foam liner and cotton wiper were disposed of. All surfaces of the abrasion tester were decontaminated between each carrier treatment using 95% ethanol and allowed to dry prior to treatment of the next carrier. Once all control and test carriers completed the “dry” abrasion cycle, they were incubated at 20-24°C (45-48% relative humidity) for a minimum of 15 minutes along with the non-wear controls. All carriers (non-wear controls and wear-exposed) were re-inoculated with 10 μL of test virus (amended with 6% FBS soil) at a minimum recoverable target density of 2.8-log_10_TCID_50_per carrier. The re-inoculum was spread over the carrier surface as previously described, and the carriers were dried at 20-24°C (45-48% RH) for a minimum of 30 minutes. The carriers then underwent a “wet” abrasion cycle. Following the application of a new, unused foam liner and sterilized cotton wiper onto the abrasion boat, the wiper was sprayed with sterile DI-H_2_O delivered via Preval sprayer from 75 ± 1cm distance for a 1-second duration. The wetted cotton wiper was reaffixed to the wear tester device and the carriers were each treated with four passes as described. A total of 12 complete abrasion cycles (six dry and six wet) and 11 re-inoculation events were performed per control and test formulation, per virus. Abrasion cycles alternated between dry and wet abrasions. The 12 wear and abrasion cycles were performed within a period spanning a minimum of 18 hours to greater than 24 hours (Table 1).

### Determination of Residual Sanitizing Activity

Upon completion of the 12^th^ and final wear and abrasion cycle, residual antiviral activity was assessed for commercial products TF-A and TF-B. Test (TF-A, TF-B), non-wear control, and wear-exposed control carriers were inoculated with 10 μL of stock virus (amended with 6% FBS soil) at a minimum recoverable target density of 4.8-log_10_TCID_50_per carrier. The inoculum was spread over the carrier surface as described previously and held for a contact time of five minutes. Carriers were neutralized by rinsing with 1mL of Letheen Broth Base (Neogen, Lansing, MI, USA) coupled with scraping to facilitate virus removal. The virus suspension was immediately by passed through a Sephadex G-10 (Sigma-Aldrich, St. Louis, MO, USA) filter column via centrifugation (5 minutes, 3500 x *g*). The additional carriers prepared for the cytotoxicity and neutralization validation assays were harvested in the same manner as described previously but were inoculated with 10 μL of 0% FBS MEM in lieu of the test virus. The neutralized carrier filtrates for the HCoV-229E and SARS-CoV-2 tests were diluted ten-fold using 0% FBS MEM and then plated onto 96-well trays seeded with the respective permissive host cell line prepared to confluency as described previously. Neutralized carrier filtrates for Influenza A (H1N1) were diluted (1:10) using Influenza Infection Medium. Each dilution was plated in replicates of six at 100 μL per well, and trays were incubated for 30 minutes at 37°C (HCoV- 229E and SARS-CoV-2) or 35°C [Influenza A (H1N1)] to facilitate virus-host cell adsorption. An additional 70 μL of maintenance medium [5% FBS MEM for HCoV-229E and SARS-CoV-2; Influenza Infection Medium for Influenza A (H1N1)] were then added to each well. Trays were incubated at 37°C or 35°C within a 5% CO_2_atmosphere for seven days and visualized regularly for the development of viral cytopathogenic effects (CPE) by inverted light microscopy under 10X magnification. Influenza A (H1N1) assay trays received an additional 20μL of fresh maintenance medium following 3 to 4 days of incubation to maintain optimal trypsin levels for infection.

Levels of recovered infectious viruses were calculated for each carrier using the Spearman-Karber algorithm as described previously. The mean virus recoveries for each quadruplicate set of control (non-abrasion and abrasion) and test carriers treated using TF-A and TF-B were determined, and the residual virucidal effectiveness for these formulations were computed relative to the mean virus recoveries from the abraded control carriers. The effects of abrasion alone on virus infectivity were assessed by comparing the mean virus recovery from the abraded control carriers to levels recovered from non-abraded control carriers.

## Results

The mean virus recovery values measured for the initial inoculum, re-inoculations, and residual antiviral activity assessments for test formulations TF-A and TF-B met the minimum log_10_TCID_50_carrier recovery requirements and are shown in Table 3. Standard deviation values for initial inoculation and re-inoculations are not provided due to the small sample size (n=2). Variability within each quadruplicate set of carriers was less than 0.50 log_10_for Non-Abrasion and Abrasion Controls across the three test viruses, and the process of wear and abrasion alone did not impact levels of inoculated infectious viruses during the subsequent five-minute contact time to determine residual activity.

**Table 3.**
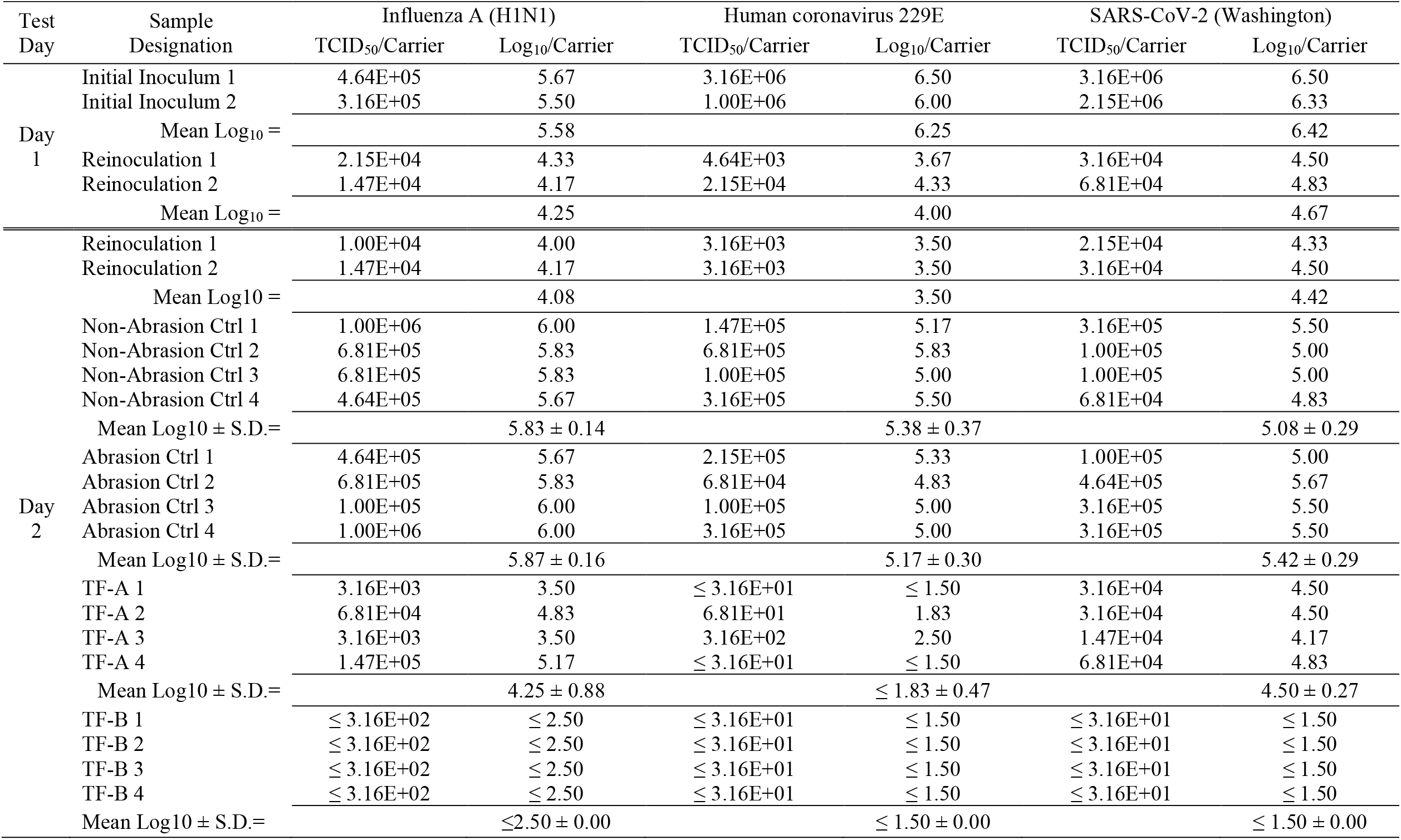
Carrier Virus Recovery Values Measured during Wear and Abrasion Procedure

**Table 4.** Log_10_Inactivation of Influenza A (H1N1), HCoV-229E, and SARS-CoV-2 following a 5-Minute Exposure to Residual Levels of Test Formulations TF-A and TF-B

| Sample Designation | Influenza A (H1N1) (Log<sub>10</sub> Reduction)<sup>a,b,c</sup> | Human coronavirus 229E (Log<sub>10</sub> Reduction)<sup>a,b,c</sup> | SARS-CoV-2, Washington (Log<sub>10</sub> Reduction)<sup>a,b,c</sup> |
| --- | --- | --- | --- |
| Non-Abrasion Control (n=4) | N.A. | N.A. | N.A. |
| Abrasion Control (n=4) | + 0.04 | 0.21 | + 0.33 |
| TF-A (n=4) | 1.62 | 3.33 | 0.92 |
| TF-B (n=4) | ≥ 3.38 | ≥ 3.67 | ≥ 3.92 |

<sup>a</sup>N.A.: Not applicable.
<sup>b</sup>Log<sub>10</sub> reductions for Abrasion Control calculated relative to Non-Abrasion Control recovery mean.
<sup>c</sup>Log<sub>10</sub> reductions for TF-A and TF-B calculated relative to Abrasion Control recovery mean.

Viable Influenza A (H1N1), HCoV-229E, and SARS-CoV-2 viruses were recovered following five minutes of contact time with residual levels of Test Formulation A (TF-A), which was most effective against HCoV-229E with 2 out of the 4 carriers demonstrating virus levels below the limit of detection (1.50 log_10_) (Table 3). In contrast, infectious Influenza A (H1N1) and SARS-CoV-2 (Washington) viruses were recovered from all (4 out of 4) residual TF-A carriers after five minutes of exposure. Although >3-log_10_infectious viruses were recovered following exposure to TF-A, analysis using the Paired Student’s t-test (p<0.05) revealed that the differences in recovery were significant for Influenza A (H1N1) (p=0.03) and SARS-CoV-2 (p=0.02) compared to their respective mean abrasion controls. This indicates that there was some antiviral activity imparted by residual TF-A that decreased infectious virus numbers; however, the degree of antiviral activity was not sufficient to achieve the 3-log_10_reduction requirement for residual sanitization (Table 3).

Influenza A (H1N1), HCoV-229E and SARS-CoV-2 were consistently inactivated to levels below the limit of detection within five minutes of contact with residual Test Formulation B (TF-B) (Tables 2 and 3). Toxicity was observed of each of the three host cell lines for carriers harvested during the residual activity assessments, as well as those designated for the cytotoxicity and neutralization validation controls. The MRC-5 and Vero E6 cell lines demonstrated cytotoxic effects in the lowest plated dilution, thereby yielding a cytotoxicity titer of 1.50 log_10_. However, no viral CPE was observed for HCoV-229E or SARS-CoV-2 beyond the level of toxicity for the carriers assessed for TF-B residual activity. Further, log_10_reductions of HCoV-229E and SARS-CoV-2 measured ≥ 3.67 log_10_and ≥ 3.92 log_10_, respectively, and sanitization was achieved within 5 minutes of contact with residual TF-B. Increased cytotoxicity differing by an order of magnitude was observed on the MDCK cell line, resulting in a higher detection limit of 2.50 log_10_. Still, no changes to host cell morphology indicative of infection by Influenza A (H1N1) were observed beyond the level of cytotoxicity, and residual sanitizing activity during the five-minute exposure to TF-B measured ≥ 3.38 log_10_.

## Discussion

Two commercially-available, ready-to-use spray formulations (coded herein as TF-A and TF-B) characterized by varying types and concentrations of quaternary ammonium actives and the presence or absence of a binder chemistry, were evaluated for continuous residual sanitizing activity on glass carriers using a wear and abrasion procedure against Influenza A (H1N1), HCoV-229E, and SARS-CoV-2 (Washington) in the presence of 6% organic soil. The actives comprising TF-A are common-use quaternary ammonium compounds. In contrast, TF-B contains a quaternary ammonium compound that carries a 24-hour US EPA residual kill claim of 99.9% against bacteria as well as a polyethyloxazoline binder. Following wear and abrasion testing TF-A performed variably, achieving greater than 3- log_10_inactivation for HCoV-229E yet less efficacy against Influenza A (H1N1) and SARS-CoV-2 (Table 3). In comparison, residual TF-B performed consistently against each of the three test viruses, achieving reductions of ≥3- log_10_to levels below detection and beyond the observed levels of cytotoxicity. Of the two formulations, TF-B rendered self-sanitizing properties to the glass carriers even after 12 alternating wet-dry cycles of wear and abrasion with repeated inoculations of test virus in the presence of a 6% (v/v) organic soil. It is important to acknowledge that the study scope was limited to enveloped viruses – two coronaviruses and one influenza virus. Extrapolation of the observed virus recovery levels and inactivation trends documented to other enveloped viral pathogens of concern as well as non-enveloped viral agents should therefore be avoided.

Inhalation of infectious aerosols is the primary mode of transmission for respiratory pathogens. SARS-CoV-2 and Influenza A have been detected in the air of hospitals and other high-traffic public spaces in locales worldwide principally using molecular methods including quantitative PCR (Xie et al., 2020; Chamseddine et al., 2021; Hadei et al., 2021; Passos et al., 2021). Infectious SARS-CoV-2 has also been detected in community-generated aerosols (Lednicky et al., 2020) and within the residential air spaces of infected persons (Vass et al., 2022). The settling of aerosols following their release by infected individuals can contaminate environmental surfaces over time within built environments. Genomic RNA of SARS-CoV-2 and Influenza has been isolated from surfaces in public spaces (Killingley et al., 2016; Fong et al., 2020; Harvey et al., 2020; da Silva et al., 2021). Detection of viable SARS-CoV-2 from surfaces has been less frequent (Marcenac et al., 2021; Vass et al., 2022), but this may be due to the abundance of studies employing molecular techniques rather than cell culture infectivity methods (Goncalves et al., 2021). Once deposited onto surfaces, decay of infectious SARS-CoV-2 and Influenza A are dependent on several environmental factors including temperature and surface type (Thompson et al., 2017; Riddell et al., 2020; Ronca et al., 2021). Still, the role of fomites in the transmission of infectious respiratory diseases has been questioned in the scientific community (Killingley et al., 2016; Goldman, 2021), and quantitative microbial risk assessment (QMRA) modeling deemed surface transmission as low-risk (<10^−6^) for contracting infectious SARS-CoV-2 (Pitol and Julian, 2021).

Nevertheless, the perception of risk by the public at-large should remain a critical factor of consideration. The COVID-19 pandemic saw an increase in the practice of protective behaviors such as disinfection by individuals who sought to protect their health and well-being (Vally, 2020). However, a survey conducted by the American Cleaning Institute in 2020 revealed that only 26% of respondents adhered to label use directions during routine household disinfection practices, with an equal percentage acknowledging that surfaces are wiped dry immediately after spraying (The American Cleaning Institute, 2020). Antimicrobial products that are formulated for enhanced adhesion to surfaces, such as Test Formulation B (TF-B) evaluated herein, present an alternative to standard disinfectants which may be easily removed by wear and abrasion (i.e., wiping) under dry and moist conditions. This becomes particularly critical as surfaces tend to become re-contaminated over time, which was simulated by the regular re-inoculations performed in the current study. While previous research efforts have demonstrated the effectiveness of self-disinfecting surfaces, most of the actives studied were embedded materials comprised, in part, of titanium dioxide, metals, or novel specialized polymers that were under development and not broadly available for commercial use (Querido et al., 2019; Ikner et al., 2021; Ikner and Gerba, 2021). Published studies employing wear and abrasion with repeated reinoculations to assess the residual efficacy of spray products are scant (Rutala et al., 2021). This paper presents antiviral effectiveness data following the use of a rigorous wear and abrasion protocol to demonstrate and compare the residual self-sanitizing capability of commercially available spray disinfectants, one with a US EPA residual kill claim against bacteria, against three respiratory pathogens of concern to public health in the presence of a 6% organic soil load.

## Statements and Declarations

All authors contributed to the study conception and design. Material preparation, data collection and analysis were performed by Luisa Ikner, Andrew Rabe, and Charles Gerba. The first draft of the manuscript was written by Luisa Ikner and Andrew Rabe and all authors commented on previous versions of the manuscript. All authors read and approved the final manuscript.

This study was funded by The Procter & Gamble Company.

Luisa A. Ikner and Andrew Rabe have no financial or non-financial interests to declare. Charles P. Gerba has received research funding from The Procter and Gamble Company.

## Notes

### Competing Interest Statement

This study was funded by The Procter & Gamble Company.
Luisa A. Ikner and Andrew B. Rabe have no financial or non-financial interests to declare. Charles P. Gerba has received research funding from The Procter and Gamble Company.

